# ColocAI: artificial intelligence approach to quantify co-localization between mass spectrometry images

**DOI:** 10.1101/758425

**Authors:** Katja Ovchinnikova, Alexander Rakhlin, Lachlan Stuart, Sergey Nikolenko, Theodore Alexandrov

**Author notes:** These authors equally contributed (Alexander Rakhlin, Lachlan Stuart).

## Abstract

**Motivation:** Imaging mass spectrometry (imaging MS) is a prominent technique for capturing distributions of molecules in tissue sections. Various computational methods for imaging MS rely on quantifying spatial correlations between ion images, referred to as co-localization. However, no comprehensive evaluation of co-localization measures has ever been performed; this leads to arbitrary choices and hinders method development.

**Results:** We present ColocAI, an artificial intelligence approach addressing this gap. With the help of 42 imaging MS experts from 9 labs, we created a gold standard of 2210 pairs of ion images ranked by their co-localization. We evaluated existing co-localization measures and developed novel measures using tf-idf and deep neural networks. The semi-supervised deep learning Pi model and the cosine score applied after median thresholding performed the best (Spearman 0.797 and 0.794 with expert rankings respectively). We illustrate these measures by inferring co-localization properties of 10273 molecules from 3685 public METASPACE datasets.

**Availability and Implementation:** https://github.com/metaspace2020/coloc

## Introduction

Metabolites and lipids play key roles in fueling and making up cells, ultimately determining their types and states. Concentrations of metabolites and lipids are carefully regulated to maintain homeostasis in tissues, organs, and organisms, and are profoundly and sometimes irreversibly altered in disease. Capturing spatial distributions of molecules in tissue sections is a prerequisite for any hypothesis-driven or discovery-oriented investigation of biology and medicine on the levels of tissues and the organism. In the past two decades, a window of opportunity has been opened by the development and further maturation of imaging mass spectrometry (imaging MS), a powerful and versatile technology for spatial molecular analysis (Doerr, 2018; Dreisewerd and Yew, 2017; Buchberger *et al.*, 2018) with a particular interest in clinical (Vaysse *et al.*, 2017) and pharmaceutical applications (Schulz *et al.*, 2018). For a tissue section, imaging MS generates a hyperspectral image encompassing thousands to millions of ion images, each image representing the distribution of a particular molecule or several molecules in the section. Rapid development and growing popularity of imaging MS, as well as the high dimensionality and sheer size of generated data, measuring up to hundreds of gigabytes for a tissue section, have stimulated the development of computational methods and software (Alexandrov, 2012). Various methods have been developed for low-dimensional data representation (based on PCA, NMF, t-SNE, bi-clustering), finding spatial regions of interest with spatial segmentation, search for markers associated with a region of interest, and, recently, for metabolite annotation (Palmer *et* al., 2017). Many of these methods use some measure of spatial similarity between ion images, often referred to as *spatial co-localization*. Various measures for quantifying co-localization have been proposed, including the Pearson correlation, cosine score, Euclidean L_2_-measures (McCombie *et al.*, 2005; McDonnell *et al.*, 2008; Alexandrov *et al.*, 2010; Alexandrov, 2012) sometimes applied to transformed images, e.g., after hotspot removal or log-transformation (Watrous *et al.*, 2011). Recently, new measures adopted from other fields have been proposed, including the Structural Similarity Index (SSIM) and hypergeometric similarity measure (Kaddi *et al.*, 2011; Ekelöf *et al.*, 2018; Aaron *et al.*, 2018). However, despite the ubiquity of using spatial co-localization in imaging MS and a variety of measures proposed, no rigorous and comprehensive evaluation of co-localization measures has ever been performed.

This leads to arbitrary and often *ad hoc* choice of a co-localization measure in every particular study, lab, or software package. Moreover, it hinders the progress of imaging MS methods since new co-localization measures are faced with scepticism without objective criteria to demonstrate their advantages. This gap has persisted for over a decade due to the lack of ground truth data that would allow one to evaluate a measure objectively. Obtaining ground truth data is challenging because it requires a comprehensive inventory of which molecules are represented in imaging MS data and which of them are co-localized. This is not possible for tissues and hardly possible even for authentic standards due to our limited understanding of ionization of complex mixtures.

Here, we are addressing this apparent gap by presenting ColocAI, an artificial intelligence approach to quantify co-localization between ion images. First, we present a gold standard set of pairs of ion images ranked by imaging MS experts by the perceived co-localization. This effort was enabled by METASPACE, the open knowledge base of spatial metabolomes (Alexandrov *et* al., 2019), through being able to select a large number of public representative datasets, employ modern web-based technologies for user-friendly and facilitated image ranking, engage a large number of experts, and consolidate their rankings into a high-quality gold standard set. Second, using the gold standard set of pairs of images manually ranked by their co-localization, we have evaluated a variety of co-localization measures, including the cosine score, Pearson correlation, and SSIM. Moreover, we propose several novel measures for co-localization, e.g., using tf-idf adopted from natural language processing as well as approaches based on deep learning.

We found the semi-supervised deep learning-based Pi model as well as the cosine score applied after median thresholding to be the most optimal spatial co-localization measures for imaging MS. We propose to use them in data analysis methods relying on co-localization. Our work provides a gold standard set (available at GitHub^1^) which can be used for evaluating future measures, and in general illustrates how artificial intelligence approaches enabled by open-access data, web technologies, community engagement, and deep learning open novel avenues to addressing long-standing challenges in imaging MS.

## Methods

### Experiment design to collect expert knowledge

In artificial intelligence and computer vision, a gold standard set is a collection of images manually tagged or ranked by experts called rankers. Having a gold standard set enables training and evaluation of machine learning models and algorithms. However, creating an unbiased, representative, and high-quality gold standard set is a substantial challenge on its own. To the best of our knowledge, there exists no gold standard set of co-localized images for imaging MS. We aimed at creating a gold standard set that would quantify the perceived by experts degree of co-localization for different ions. We designed the gold standard set to consist of *target-comparison sets* where each set includes one *target ion* and 10 *comparison ions* ranked according to their co-localization with the target ion.

To create a gold standard set of co-localized ion images, we selected public datasets from METASPACE with the aim to have a manageable number of diverse yet representative high-quality datasets from different labs. First, we selected labs with at least three active contributors of public data, 9 labs in total. For every lab, we selected active contributors to METASPACE, 42 rankers in total. We aimed at asking each ranker to rank up to 20 sets. For each lab, we randomly selected round(20 * N_TL * 2 / 3) public datasets submitted by this lab to METASPACE, where N_TL is the number of rankers from a given lab.

From each dataset, we randomly selected one target ion and 10 comparison ions constituting a target-comparison set. We then used the RankColoc web app (described later) to go through the target-comparison sets and exclude noisy images or images with only a few pixels. For each lab, we aimed at obtaining round(20 * N_TL / 3) high quality sets, although it was not always possible due to the quality of the datasets. This allowed us to have the same target-comparison set ranked by three rankers to test for the ranking consistency and to obtain average ranks.

### Pilot study

Before creating the gold standard set, we ran a pilot study to investigate the difficulty of ranking ion images in the target-comparison sets according to their co-localization, as well as to learn potential pitfalls and obstacles of the ranking process. The pilot study was basically a full study including dataset selection, web app implementation, rankers recruitment, gold standard set creation, and agreement evaluation, but performed in a smaller format with 5 rankers only.

### Web app for manual ranking of ion images

The RankColoc web app^2^ was developed with the aim to facilitate image ranking as well as help inspect ranked sets. For a public dataset in METASPACE, the web app downloads ion images from METASPACE using the GraphQL API^3^, and shows a target and 10 comparison ion images. The web app helps a ranker rank each comparison image from 0 to 9 by dragging and dropping it into one of the ten rank boxes or leave it unranked. Several images can be assigned the same rank. The web app page includes instructions for rankers. For each ranker, we assigned a collection of target-comparison sets and generated unique URLs containing the sets and the ranker ID. Ranking results were stored in real time, associated with the ranker ID, and could be opened by either the same ranker or a curator. Figure 1 shows screenshots of RankColoc web app with examples of the ranked sets. Supplementary Video 1 illustrates the ranking process.

**Figure 1.**
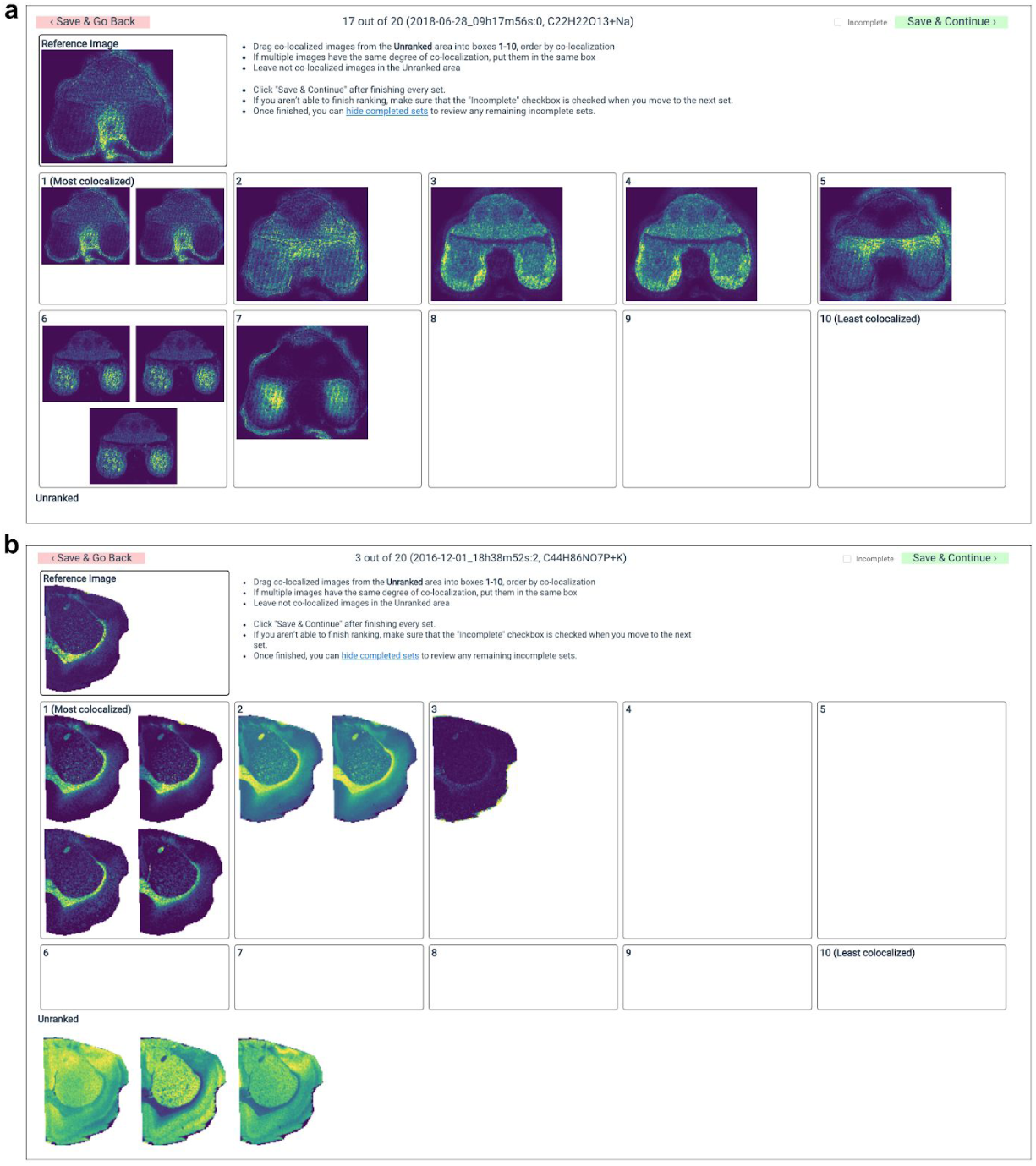
Screenshots of the RankColoc web app showing two target-comparison sets ranked by experts. a. MALDI-imaging dataset from a wheat seed section, submitted to METASPACE by Dhaka Bhandari, Justus Liebig University Giessen. b. MALDI-imaging dataset from a rat brain tissue section, submitted to METASPACE by Berin Boughton, University of Melbourne^4^.

### Evaluating obtained rankings

We assessed the complexity of the task and reproducibility of the rankers’ judgements by calculating pairwise correlations between the rankers, i.e., correlations between their ranks of comparison ion images in the same sets. The images left unranked (i.e., perceived by rankers as completely not co-localized with the target ion image) were assigned the rank 10. We computed average Spearman and Kendall ranker-pairwise rank correlation for each lab, ranker, and set.

### Creating the gold standard set

To ensure the high quality of the resulting gold standard set, we have excluded (a) sets for which the average Spearman pairwise correlation between rankers was less than 0.4, and (b) rankers whose average Spearman correlation with other rankers was less than 0.4. After some rankers were excluded, some sets ended up with just one ranking. We excluded those sets as well.

The resulting gold standard set contains pairs of ion images (target image and a comparison image) with each pair assigned an average rank across three rankers. The ranks range from 0, representing the highest co-localization, to 10, representing the lowest or no co-localization.

### Co-localization measures

Our implementation of the co-localization measures is available at the GitHub repository ^5^.

#### Measures that do not require learning

##### Correlation and cosine-based measures

First, we considered the commonly-used co-localization measures: Pearson correlation, Spearman correlation, and cosine similarity applied to flattened ion images, i.e., one-dimensional vectors of pixel intensities.

##### Structural similarity (SSIM) measure

Following Ekelöf *et al.*, 2018), we considered the structural similarity (SSIM) index (Wang *et al.*, 2004) with the Gaussian weights.

##### Tfidf-based measure

We developed a measure based on the term frequency–inverse document frequency (tf-idf) concept from the field of natural language processing (Leskovec *et al.*, 2014). Using flattened ion images, we calculated the tf-idf value for each pixel-ion pair to quantify how important a pixel *p* is for the particular ion *i* with respect to all ions in the dataset *D* :

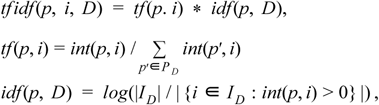

where *P*_*D*_ is the set of all pixels in *D, I_D_* is the set of all ions in *D*, and *int*(*p, i*) is the intensity of *i* in *p*. We then created tf-idf vectors of the same dimensionality as the intensity vectors and quantified co-localization of ion images as the cosine similarity between the corresponding tf-idf vectors.

##### Image transformations

For all considered ion intensity-based measures, we applied the following transformations to the ion image prior to calculating co-localization: 1) removing hotspots with intensities of greater than 0.99 quantile; 2) applying the median filter with a square window of size ranging from 1 (no filter applied) to 5 with step 1; 3) applying quantile thresholding, namely filtering out pixels with intensities below a quantile value for quantiles ranging from 0 to 0.9 with step 0.05. Evaluation whether using a transformation is beneficial as well as optimizing the size of the median filter and the quantile value was performed using the 5-fold cross-validation for each measure. Measures with the best performing filters were then applied to the entire gold standard set.

#### Measures based on deep learning

With the advent of deep learning in artificial intelligence, models based on neural networks have become the method of choice for processing unstructured data such as images. Therefore, in our study we have developed several methods exploiting current state of the art deep learning approaches that would learn ion co-localization from the gold standard set.

##### Xception-based model

This model, illustrated in Figure 2, is based on the well-known Xception convolutional architecture designed to extract informative features from images (Chollet, 2017). We introduced the following modifications. First, the input has 2 channels corresponding to the target and comparison ion images. The 2 channels pass through the Xception architecture without the final classification layer, which in our case is replaced with a regression output. The Xception-based model is supervised, and its target variable is the rank as specified in the gold standard set, with the Mean Squared Error (MSE) loss function.

**Figure 2.**
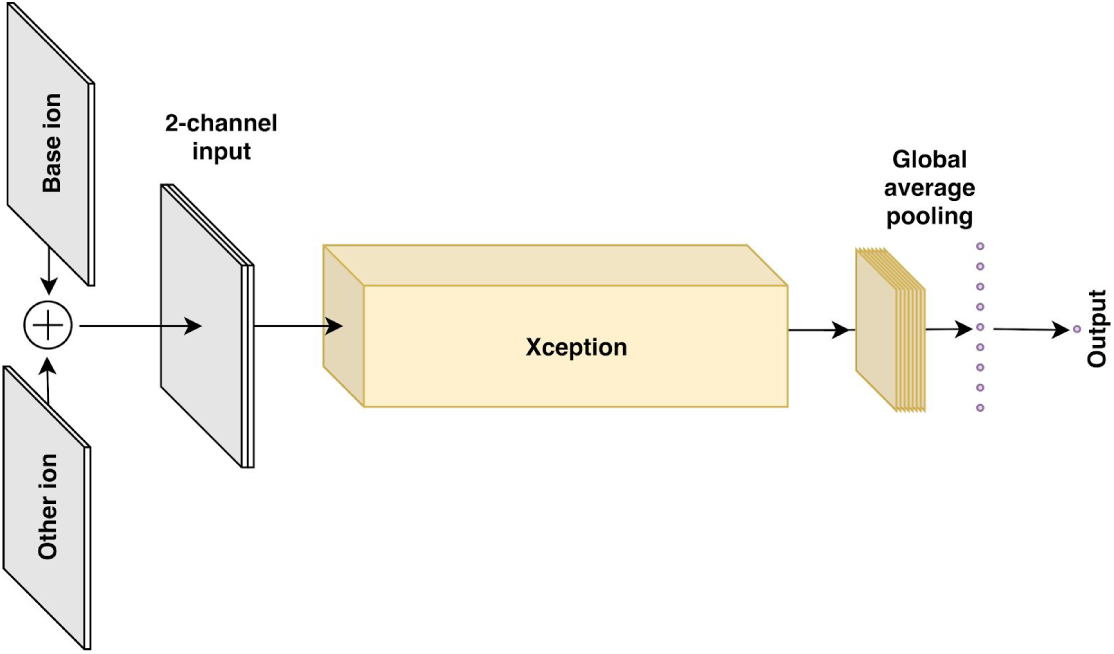
Architecture of the Xception-based deep learning model.

##### Mu model

The Mu model is a variation of the Xception-based model with the difference that the top layers are replaced with two 2048-dimensional outputs followed by a discriminator. The mu model encodes a pair of ion images into two 2048-dimensional representations, computed image similarity as the Pearson correlation coefficient between the representations, and then regresses the similarity score onto the rank target with the MSE loss.

##### Unsupervised UMAP

We developed a model based on the Uniform Manifold Approximation and Projection (UMAP), a recently developed nonlinear dimensionality reduction technique with broad applications in biology (McInnes *et al.*, 2018). In this model, we applied UMAP to embed flattened ion images (i.e., one-dimensional vectors of pixel intensities) into 20-dimensional space using the “cosine” distance metric. After the unsupervised embedding model defined the distance between ion images, we calculated the Pearson correlation coefficient between the corresponding embedded vectors to rank comparison images with respect to the target image. This model is unsupervised and does not use the gold standard set.

##### UMAP+GBT model

Since in our case supervision is actually possible, we extended the UMAP model with a supervised model on top. Namely, we used gradient boosted trees (GBT), a state of the art regression model (Chen and Guestrin, 2016), feeding UMAP 20-dimensional features as input and regressing them onto rank targets from the gold standard set with the MSE loss function.

##### Pi model

The pi model is based on the recently developed approach of temporal ensembling for semi-supervised learning (Laine and Aila, 2016). This approach uses an ensemble of network outputs from different training epochs as quasi-targets for training on unlabeled samples, which has been shown to significantly improve the final model quality. In our case, the pi model follows the general architecture of the Xception-based model, but the last layers are replaced with two heads for two loss components:

- supervised loss as in the Xception-based model, the MSE between the network prediction and the rank;
- unsupervised loss intended to stabilize the network prediction; we define it as the squared error between network predictions on a pair of ion images and the same pair of ion images subjected to various image augmentations (intensities and geometric transformations).

The unsupervised loss component has allowed us to use ~40,000 unlabelled ion images from 3685 public METASPACE datasets, gathering ~56,000 unlabelled pairs from them for training in addition to the labeled gold standard set.

### Evaluation of the co-localization measures

For each set, we calculate Spearman and Kendall correlation coefficients separately for each target-comparison set and report the mean and median values ovell all sets.

## Results

### The co-localization gold standard set

We have initially selected 239 datasets from METASPACE with 304 target-comparison sets of ion images for 42 rankers from 9 labs (Table 1).

**Table 1.** Information about the co-localization gold standard set created from public METASPACE datasets contributed by 9 labs with target-comparison sets ranked manually by 42 experts from these labs. The final version was obtained after filtering out rankers and sets with agreement less than 0.4 and sets ranked by one ranker only.

| Lab | Number of rankers | Number of datasets | Number of sets | Average pairwise agreement |  |  |  |
| --- | --- | --- | --- | --- | --- | --- | --- |
|  |  |  |  | Spearman |  | Kendall |  |
|  |  |  |  | mean | median | mean | median |
| University of Copenhagen | 8 | 66 | 78 | 0.534 | 0.730 | 0.489 | 0.620 |
| EMBL | 8 | 33 | 53 | 0.763 | 0.804 | 0.666 | 0.710 |
| University of Melbourne | 6 | 29 | 40 | 0.806 | 0.836 | 0.732 | 0.742 |
| JLU Giessen | 5 | 29 | 33 | 0.612 | 0.688 | 0.545 | 0.597 |
| IBMP | 3 | 20 | 20 | 0.638 | 0.681 | 0.550 | 0.604 |
| PNNL | 3 | 20 | 20 | 0.686 | 0.687 | 0.596 | 0.609 |
| MPI Bremen | 3 | 19 | 20 | 0.843 | 0.925 | 0.787 | 0.879 |
| UT Austin | 3 | 16 | 20 | 0.808 | 0.866 | 0.745 | 0.770 |
| M4I | 3 | 7 | 20 | 0.730 | 0.680 | 0.660 | 0.608 |
| Total | 42 | 239 | 304 | 0.700 | 0.773 | 0.629 | 0.666 |
| Final version after filtering out rankers and sets | 38 | 182 | 234 | 0.791 | 0.800 | 0.711 | 0.708 |

### Pilot study

The pilot study was crucial to inform us about the complexity and subjectivity of the task and to design the final version of the web app and the study accordingly. In particular, we learned that ranking comparison images was more natural for the rankers than ordering them because this allowed the rankers to assign the same rank to several comparison images. Selecting high-quality datasets and skipping noisy ion images and images with just a few non-zero pixels was crucial for obtaining reproducible rankings. Some rankers preferred to leave non-co-localized images unranked and we have implemented this option for the final study.

### Agreement between experts

Table 1 shows the average pairwise ranker correlation values for each lab that represent agreement between rankers. Note that the agreement values cannot and shall not be compared across different labs because every lab ranked images from different METASPACE datasets. For example, ranking images with a simple and clear spatial structure led to higher agreement values. The mean correlation across all sets was 0.700 (Spearman) and 0.629 (Kendall). After excluding sets and rankers with low agreement, the mean agreements for the final version of the gold standard set is 0.791 (Spearman) and 0.711 (Kendall).

### The gold standard set

The final version of the gold standard set includes 234 sets with 2210 ion image pairs from 182 public imaging datasets from METASPACE ranked by 38 rankers from 9 labs, available at https://github.com/metaspace2020/coloc/tree/master/GS. The datasets represented human (37%), mouse (21%), pig (7%), rat (6%) and other organisms; brain (27%), kidney (11%), skin (9%), seed (4%) and other organs; MALDI (84%) and DESI (16%) ionisation; DHB (44%), DAN (17%), DHA (6%), BPYN (5%) and other MALDI matrices; Orbitrap (69%) and FTICR (31%) mass analyzers; positive (68%) and negative (32%) polarity. For every target-comparison pair of ion images, average rank across three rankers has been assigned. The ranks range from 0, representing the highest co-localization, to 10, representing the lowest or no co-localization.

### Evaluation of co-localization measures

#### Measures that do not require learning

Table 2 shows the performance of co-localization measures requiring no learning, measured using the gold standard set as Spearman and Kendall rank correlation to the expert rankings. For each measure, we show its best performance and optimal image transformation parameters. The best performing measure is the cosine similarity with quantile threshold 0.5, without hotspot removal and with median filter with window size 3. The second best measure is the Pearson correlation measure with no image transformation applied. The SSIM measure recently proposed in the context of imaging MS (Ekelöf *et al.*, 2018) was outperformed by other measures.

**Table 2.** Performance of the co-localization measures requiring no learning in terms of Spearman and Kendall correlation coefficients with the optimal parameters for image transformations.

| Co-localization measure | Correlation with the expert rankings in the gold standard set |  |  |  | Optimal parameters of image transformations |  |  |
| --- | --- | --- | --- | --- | --- | --- | --- |
|  | Spearman |  | Kendall |  | Quantile for thresholding | Hotspot removal | Median window size |
|  | mean | median | mean | median |  |  |  |
| Cosine | **0.794** | **0.849** | **0.682** | **0.720** | **0.5** | **No** | **3** |
| Tfidf-cosine | 0.769 | 0.825 | 0.653 | 0.689 | 0.65 | No | 0 |
| Spearman | 0.737 | 0.783 | 0.620 | 0.647 | 0.7 | No | 0 |
| Pearson | 0.788 | 0.838 | 0.674 | 0.719 | 0 | No | 0 |
| SSIM | 0.559 | 0.623 | 0.449 | 0.488 | 0 | No | 0 |

Table 3 shows the effect of using different types of image transformation on the performance of the cosine measure. Surprisingly, applying hotspot removal did not improve the performance. Denoising images by using the moving median filter improves the performance only marginally (Spearman correlation from 0.792 to 0.794), whereas using quantile thresholding led to a significant improvement (Spearman correlation from 0.779 to 0.794).

**Table 3.** The effect of using different types of image transformations onto the performance of the cosine-based measure.

| Image transformation applied before calculating cosine similarity | Correlation with the expert rankings in the gold standard set |  |  |  | Optimal parameters of image transformations |  |  |
| --- | --- | --- | --- | --- | --- | --- | --- |
|  | Spearman |  | Kendall |  | Quantile value | Hotspot removal | Median window size |
|  | mean | median | mean | median |  |  |  |
| Quantile thresholding, hotspot removal, denoising | **0.794** | **0.849** | **0.682** | **0.720** | **0.5** | **No** | **3** |
| Quantile thresholding, hotspot removal (no denoising) | 0.792 | 0.844 | 0.681 | 0.720 | 0.5 | No | - |
| Hotspot removal, denoising (no quantile thresholding) | 0.779 | 0.842 | 0.665 | 0.719 | - | No | 3 |

#### Measures based on deep learning

Table 4 shows the performance of co-localization measures based on deep learning models. The Pi model achieved the best performance, with a slight improvement over cosine similarity and nearly reaching the human-to-human agreement between the experts in our study. This is no surprise since the Pi model is a state-of-the-art semi-supervised model that makes use of both labeled data from the gold standard set and unlabeled data from METASPACE. What is interesting is that the purely unsupervised UMAP model performed on par with supervised Xception and UMAP+GBT, its own version enhanced with supervision through gradient boosted trees, and outperformed the Mu model. This may indicate that the structural properties of co-localization are relatively evident in the data, and once these properties are extracted the rest is just “noise”. This would imply that our results are already close to perfect given the “noise” inherent in the notion of co-localization, and would also explain why the best model results are so close to the human-to-human agreement.

**Table 4.** The performance of deep learning-based models measured as Spearman and Kendall correlations with expert rankings in the gold standard set. The best performance is achieved by the semi-supervised Pi model that makes use of both labelled and unlabelled data.

| Deep Learning-based model | Correlation with the expert rankings in the gold standard set |  |  |  |
| --- | --- | --- | --- | --- |
|  | Spearman |  | Kendall |  |
|  | mean | median | mean | median |
| Xception model | 0.777 | 0.820 | 0.682 | 0.716 |
| **Pi model** | **0.797** | **0.847** | **0.712** | **0.752** |
| Unsupervised UMAP | 0.761 | 0.827 | 0.656 | 0.686 |
| UMAP+GBT | 0.758 | 0.845 | 0.672 | 0.741 |
| Mu model | 0.725 | 0.804 | 0.638 | 0.705 |

### Inferring molecular relationships by mining public METASPACE data

We have applied the best derived co-localization measures to illustrate how they can be used on a large scale to mine data from the public knowledge base METASPACE. More specifically, we aimed to infer co-localization relationships between all molecules represented in public METASPACE annotations. For this, we downloaded ion images for all annotations from 3685 public datasets in METASPACE using its API^6^, calculated co-localizations between all ion images within a dataset using either the cosine score after 0.5 quantile thresholding or deep learning Pi-model, the best performing methods in their respective classes, and averaged co-localization across all datasets. Finally, we visualized all 10273 resulting molecular formulas in a two-dimensional space using UMAP with the average co-localization used as the pre-computed distance. The annotations that are more co-localized on average are shown closer to each other (Figure 3).

**Figure 3.**
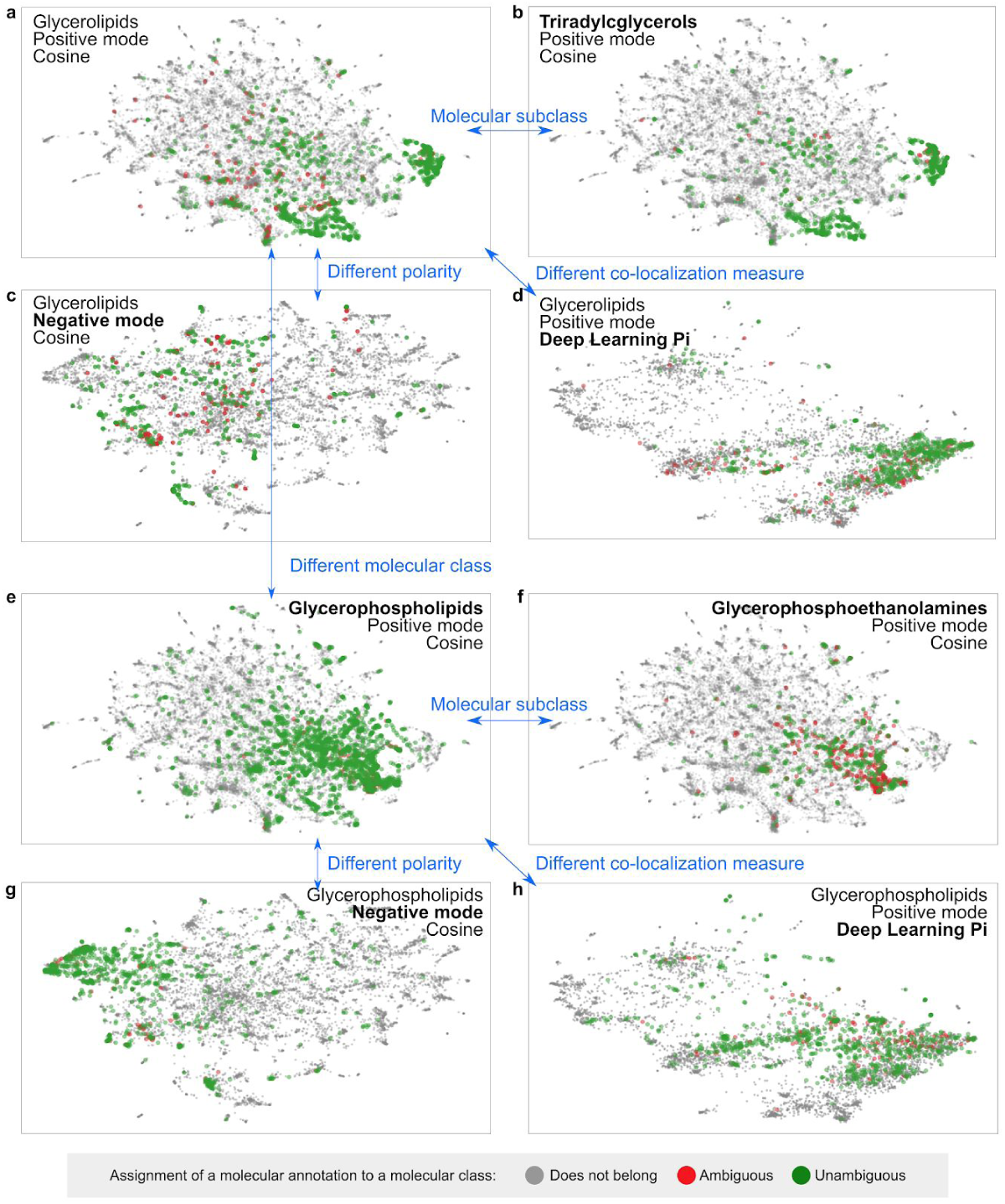
Visualization of co-localization molecular relationships as learned from METASPACE. Dots representing annotations (each corresponding to one of 10273 unique molecular formulas) are mapped based on their average co-localization across 3685 public METASPACE datasets. For a molecular class, the green color represents unambiguous assignment when all isomers belong to the class whereas the red color represents ambiguous assignment when some isotopes belong to another class.

To investigate whether the inferred co-localization properties are associated with chemical properties of the molecules, we highlighted glycerolipids, an important class of lipids which are known to be easily detectable by imaging mass spectrometry (Figure 3a). Note that the assignment of a molecular annotation (formula) to a molecular class was performed accounting for potential ambiguity, with unambiguously assigned annotations shown in green and ambiguously assigned annotations shown in red (Figure 3). One can see that glycerolipids indeed form dense clusters that indicates their high average co-localization. A subclass of glycerolipids, triradylcglycerols (with the classes names as in HDMB) represent the majority of the glycerolipids in METASPACE and form the densest clusters (Figure 3b). Sparser representation of glycerolipids in the negative polarity data (Figure 3c) illustrates the common knowledge of the positive mode being the preferred way of ionization for this class of lipids. Using another co-localization measure (deep learning-based Pi model instead of the cosine) also confirms the findings but shows a visible difference in data organization. This reflects the robust capacity of both measures to capture chemically-associated co-localization but also shows the differences between them, which can be potentially used in the future to further improve the results.

Another class of lipids, glycerophospholipids, represents a large part of molecules in METASPACE, clearly forming a cluster in the UMAP chemical space (Figure 3e). Performing examination in a way similar to Figures 3a-d, one can see that the molecular subclass of glycerophospholipids, glycerophosphoethanolamines, represents the core of the cluster of co-localized glycerophospholipids (Figure 3f). Opposite to glycerolipids, glycerophospholipids are known to be ionizable in both positive and negative modes, and this is reflected in their strong presence as well as clustered appearance on Figure 3g. Examining deep learning Pi score-based mapping (Figure 3h), one can see that despite dense spacing, there is clearly less separation visible to the class of glycerolipids (Figure 3h vs Figure 3d) compared to the cosine score-based UMAP visualization (Figure 3g vs Figure 3a), which makes cosine score-based results easier for interpretation.

## Discussion

### Gold standard

Creating a high-quality gold standard set of expert-ranked pairs of target-comparison images was possibly the most challenging part of the study. Not only it required scientific formulation of the co-localization problem and development of an experiment design able to capture the perceived extent of co-localization from the experts, but it was also the most time-consuming part of our study to organize the whole ranking experiment by selecting datasets, recruiting almost 50 experts, communicating with them, reminding them to complete the task, and when necessary coming back to them with requests for corrections. Altogether it required 95 emails solely for communicating with the rankers. Despite having expertise in performing crowdsourcing studies in imaging MS (Ovchinnikova *et al.*, 2019; Palmer *et al.*, 2015) and overwhelmingly positive support of METASPACE users in performing the ranking, running this study would not be possible without access to diverse public data in METASPACE and without using modern web technologies employed for the RankColoc web app that both critically facilitated the process. The achieved average pairwise correlation between the rankers (mean Spearman 0.791) confirms a strong inter-ranker agreement. This indicates that there is a consensus between experts with respect to perceived co-localization and, importantly, that this consensus was successfully captured in the gold standard set, thus validating our efforts.

Performing the pilot study was essential to avoid pitfalls and refine the experimental design and the web app for more objective ranking before engaging a large number of experts. Nevertheless, after performing the complete study, we see opportunities for next-level improvement. For example, in the spirit of active learning, we could choose comparison ions not randomly but those ions where our models are most uncertain about their ranking.

Taking into account the efforts necessary for producing such a gold standard set, we do not expect it to be repeated on a larger scale in the near future. However, we are considering to implement an online approach where a target-comparison set or a reduced version of it will be occasionally shown to METASPACE users. This approach would provide a continuous population of the gold standard set. However, it should be carefully designed to ensure the quality and check for consistency, since the ranking task will be split into small subtasks and performed over a period of time by a larger diverse crowd of rankers.

### Co-localization measures

Comparing the performance of the evaluated measures of co-localization (mean Spearman 0.797 for the deep learning Pi model and 0.794 for cosine similarity after median filtering) with average pairwise agreement between the rankers (mean Spearman 0.791), we suggest that the best measures approach the theoretically best performance. The slight positive difference (0.797 vs 0.791) can be due to the averaging of rankings in the produced gold standard set, thus introducing positive effects of averaging compared to the values used for the rankers agreement calculation. The fact that we potentially reached the best possible performance explains only a slight improvement when using an advanced deep learning-based method compared to cosine similarity. Another reason can be due to the specifics of the considered problem: ion images from the same dataset have the same size and structure and can be compared pixel-by-pixel after flattening; they also do not undergo changes in the view angle or brightness or other non-linear deformations that would apply to, e.g., photos used in computer vision where deep learning significantly benefits from its capacity to extract abstract visual features thus allowing comparison of different images showing the same object. Here, future efforts can be focused on developing next-level methods for spatial association between molecules that would consider “molecular microenvironment” rather than “tissue section” context.

Another effect that suggests that the considered problem is relatively simple is the comparable performance of both unsupervised and supervised methods. Another reason for this effect could be the small size of the training set (2340 pairs of ranked images); this is also supported by the best performance of a semi-supervised model which used all public METASPACE data for deriving a representation of ion images. We hope that these results, comparing a variety of deep learning models, and this discussion will be helpful for future deep learning applications in imaging MS.

Comparing the best co-localization measures (deep learning Pi model and cosine similarity after median thresholding), we investigated how well they correspond to expert ranking for each target-comparison pair from the gold standard. Figure 4 shows that there is no visible difference between these two measures: for 50% of all target-agreement sets both measures achieve high performance (Spearman correlation with the expert ranking is greater than 0.8). Despite the fact that error analysis of low-valued sets has not revealed any factors that would allow us to improve the measures, one can potentially combine the considered measures and thus achieve a better performance with an ensemble ranking. Interestingly, Figure 4 highlights that the target-comparison pairs for which both measures performed well also have visibly high values of the rankers agreement. This provides another confirmation that the developed co-localization measures reproduce the perceived co-localization when experts themselves agree on it.

**Figure 4.**
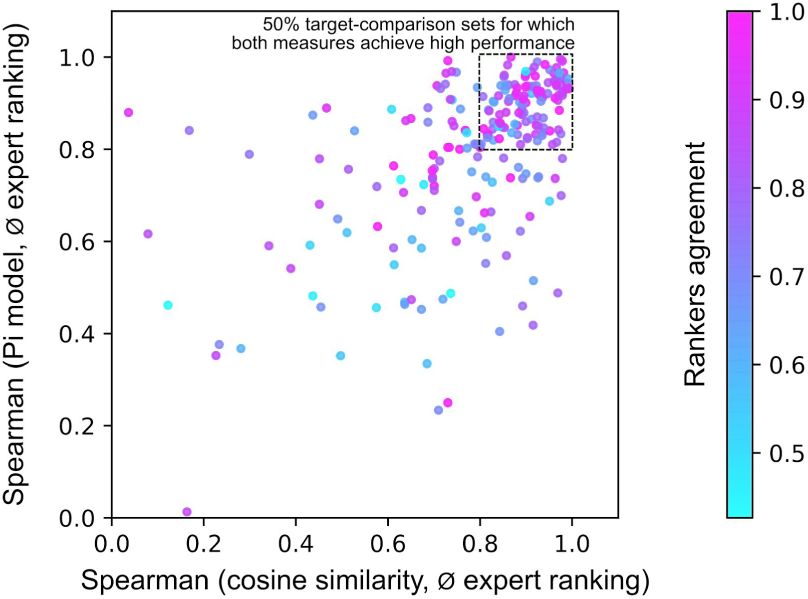
Scatterplot showing for each of 2340 target-comparison sets from the gold standard how well the best co-localization measures (deep learning Pi model, cosine similarity after median thresholding) reproduce the average expert ranking, as measured by the Spearman correlation. Each dot is colored according to the rankers agreement for the respective target-comparison set.

### Applications

A wide coverage of organisms, organs, ionization types, MALDI matrices, and mass analyzers represented in the imaging MS datasets used in the gold standard set ensures broad applicability for the findings and measures developed in this study. We expect key applications of the developed and evaluated co-localization methods to be in the search for molecular biomarkers associated with either a particular molecule or a region of interest. They should also improve distance-based methods for data analysis, e.g., representation of the full dataset using clustering of ion images (Alexandrov *et al.*, 2013). Moreover, we expect this work to provide a scientifically rigorous justification for using these measures in systems biology approaches aimed at uncovering molecular relationships between molecules by assuming the tissue representing cells of different phenotypes. Here, cutting-edge methods relying on distance or similarity measures, such as UMAP demonstrated in this paper, can replace more conventional methods such as PCA, NMF or t-SNE.

## Acknowledgements

We thank the rankers of public METASPACE data who shared their data publicly and helped us create the gold standard set: Asta Maria Joensen, Charlotte Bagger, Julian Schneemann, Meghan Friis, Fernanda Endringer Pinto, Anne Mette Handler, Andreas Danielsen, Katharina Clitherow, Sophie Jacobsen (Janfelt lab, University of Copenhagen), Mohammed Shahraz, Don Nguyen, Sergio Triana, Veronika Saharuka, Luca Rappez, Cristina Gonzalez Lopez, Aslihan Inal (EMBL), Dinaiz Thinagaran, Sanuli Paralkar, Farheen Farzana, Edita Ritmejeryte, Nicholas Sing, Berin Boughton (Metabolomics Australia, University of Melbourne), Michael Waletzko, Dhaka Bhandari, Domenic Dreisbach, Patrik Kadesch (Spengler lab, Justus Liebig University Giessen), Elisa Ruhland, Julien Delecolle, Claire Villette (Heintz lab, IBMP, University of Strasbourg), Dusan Velickovic, Christopher Anderton, Arunima Bhattacharjee (Anderton lab, Pacific Northwest National Laboratory), Manuel Liebeke, Benedikt Geier, Emilia Sogen (Liebeke lab, Max Planck Institute for Marine Microbiology Bremen), Marta Sans, Jialing Zhang, Kyana Garza (Eberlin lab, University of Texas Austin), and Shane Ellis, Pieter Kooijman, Lennart Huizing (M4I Institute, Maastricht University). This work was supported by the European Union’s Horizon 2020 program under the grant agreements 634402, 777222 (K.O., L.S., T.A.), by the Russian Foundation for Basic Research grant no. 18-54-74005 (S.N., T.A.) and the European Research Council Consolidator grant METACELL (T.A.).

## Footnotes

1 https://github.com/metaspace2020/coloc

2 https://github.com/metaspace2020/coloc/tree/master/RankColoc

3 https://github.com/metaspace2020/metaspace/tree/master/metaspace/python-client

4 METASPACE URLs for example datasets: https://metaspace2020.eu/annotations?ds=2018-06-28_09h17m56s, https://metaspace2020.eu/annotations?ds=2016-12-01_18h38m52s

5 https://github.com/metaspace2020/coloc/tree/master/measures

6 https://github.com/metaspace2020/metaspace/tree/master/metaspace/python-client

## Notes

https://github.com/metaspace2020/coloc

